# Gene regulatory network reconstruction incorporating 3D chromosomal architecture reveals key transcription factors and DNA elements driving neural lineage commitment

**DOI:** 10.1101/303842

**Authors:** Valeriya Malysheva, Marco Antonio Mendoza-Parra, Matthias Blum, Mikhail Spivakov, Hinrich Gronemeyer

## Abstract

Lineage commitment is a fundamental process that enables the morphogenesis of multicellular organisms from a single pluripotent cell. While many genes involved in the commitment to specific lineages are known, the logic of their joint action is incompletely understood, and predicting the effects of genetic perturbations on lineage commitment is still challenging. Here, we devised a gene regulatory network analysis approach, GRN-loop, to identify key *cis*-regulatory DNA elements and transcription factors that drive lineage commitment. GRN-loop is based on signal propagation and combines transcription factor binding data with the temporal profiles of gene expression, chromatin state and 3D chromosomal architecture. Applying GRN-loop to a model of morphogen-induced early neural lineage commitment, we discovered a set of driver transcription factors and enhancers, some of them validated in recent data and others hitherto unknown. Our work provides the basis for an integrated understanding of neural lineage commitment, and demonstrates the potential of gene regulatory network analyses informed by 3D chromatin architecture to uncover the key genes and regulatory elements driving developmental processes.

## Introduction

Lineage commitment is driven by a complex interplay of extrinsic and intrinsic signals acting jointly to promote global changes in gene expression and chromatin state. While many factors involved in this process have been characterised individually in a number of lineages, the mechanisms of their joint action still remain incompletely understood, hindering our understanding of normal and aberrant morphogenesis.

*Cis*-regulatory elements (CREs) on the DNA are the key “integration units” of developmental gene control. CREs recruit combinations of transcription factors (TFs), including intrinsic TFs and those acting downstream of signalling cascades. TF recruitment to CREs promotes the expression of genes that these elements physically contact through 3D chromosomal interactions (Heinz et al. 2015). CREs thereby hold the clues to the principal regulatory inputs received by developmental genes and to the logic of their expression control. However, the complexity of the regulatory landscape, including the multitudes of TFs recruited to each CRE and multiple CREs associated with most developmental genes, as well as redundancy in developmental gene control mechanisms (Cooke et al. 1997; Kafri, Levy, and Pilpel 2006; Osterwalder et al. 2018), prompts for formal analysis methods for extracting this information from relevant biological data.

Gene regulatory networks (GRNs) provide a framework to integrate information on causal relationships between multiple regulators and follow the propagation of regulatory signals in the course of a dynamic biological process, such as lineage commitment. To date, GRN analysis has been used successfully by ourselves and others to pinpoint factors and pathways underpinning normal development (Viiri et al. 2019; M.-A. Mendoza-Parra et al. 2016), cancer progression (Malysheva et al. 2016; Costa, Boroni, and Soares 2018; Verfaillie et al. 2015) and transdifferentiation (M.-A. Mendoza-Parra et al. 2016; Ieda et al. 2010). In particular, a signal propagation-based framework that we have recently developed, TETRAMER (Cholley et al. 2018; M.-A. Mendoza-Parra et al. 2016), enables the analysis of very large networks derived from genome-scale data to delineate driver nodes and paths.

Here, we present a novel signal propagation-based GRN reconstruction and analysis approach, GRN-loop, that capitalises on CREs as the pivotal links between TFs and their target genes to uncover the role of these elements and TFs recruited to them in driving gene expression changes in dynamic biological processes. GRN-loop integrates TF binding data with the temporal profiles of the transcriptome, chromatin state and 3D chromosomal architecture to capture the dynamic patterns of CRE activity and their relationships with target genes. Applying GRN-loop to a model of retinoic acid (RA)–induced early neurogenesis in P19 mouse embryonic stem cell-like cells (M.-A. Mendoza-Parra et al. 2016), we reveal key CREs and TFs driving neural lineage commitment, some of which were independently validated in recent studies and others hitherto unknown.

## Results

### Capturing the dynamics of chromosomal interactions during morphogen-induced neural commitment for GRN reconstruction

To capture the dynamics of chromosomal organization upon morphogen-induced neural lineage commitment, we performed Hi-C in P19 cells treated with retinoic acid (RA) at 0, 6 and 48 hours post-treatment. First, to detect global changes we defined topologically associated domains (TADs) using a standard method (Dixon et al. 2012), and compared the locations of TAD borders across timepoints. While a majority of TADs (~60%) were preserved during lineage commitment, in line with previous observations (Dixon et al. 2012, 2015; Nora et al. 2012), the remaining TADs showed changes in the locations of one or both borders in at least one condition (Figs. 1A, 1B, S1). We next detected individual chromatin interactions using Fit-Hi-C (Ay, Bailey, and Noble 2014) and followed the “fate” of the identified interactions across the timepoints, revealing multitudes of rewired and preserved contacts (Figs. 1C, 1D). For the purposes of GRN reconstruction, we focused on 30,332 chromosomal interactions involving the promoters of genes showing differential expression along the same time course (DEG interactions) based on our previously published transcriptome data (M. A. Mendoza-Parra et al. 2014; M.-A. Mendoza-Parra et al. 2016). A total of 16,961 DEG interactions were detected in untreated cells, with 5,388 (32%) of them maintained across all three timepoints; the remaining 13,371 DEG interactions were selectively detected in RA-treated cells (Fig. 1E). DEG interactions were used as inputs for the GRN model, as described below.

**Fig. 1.**
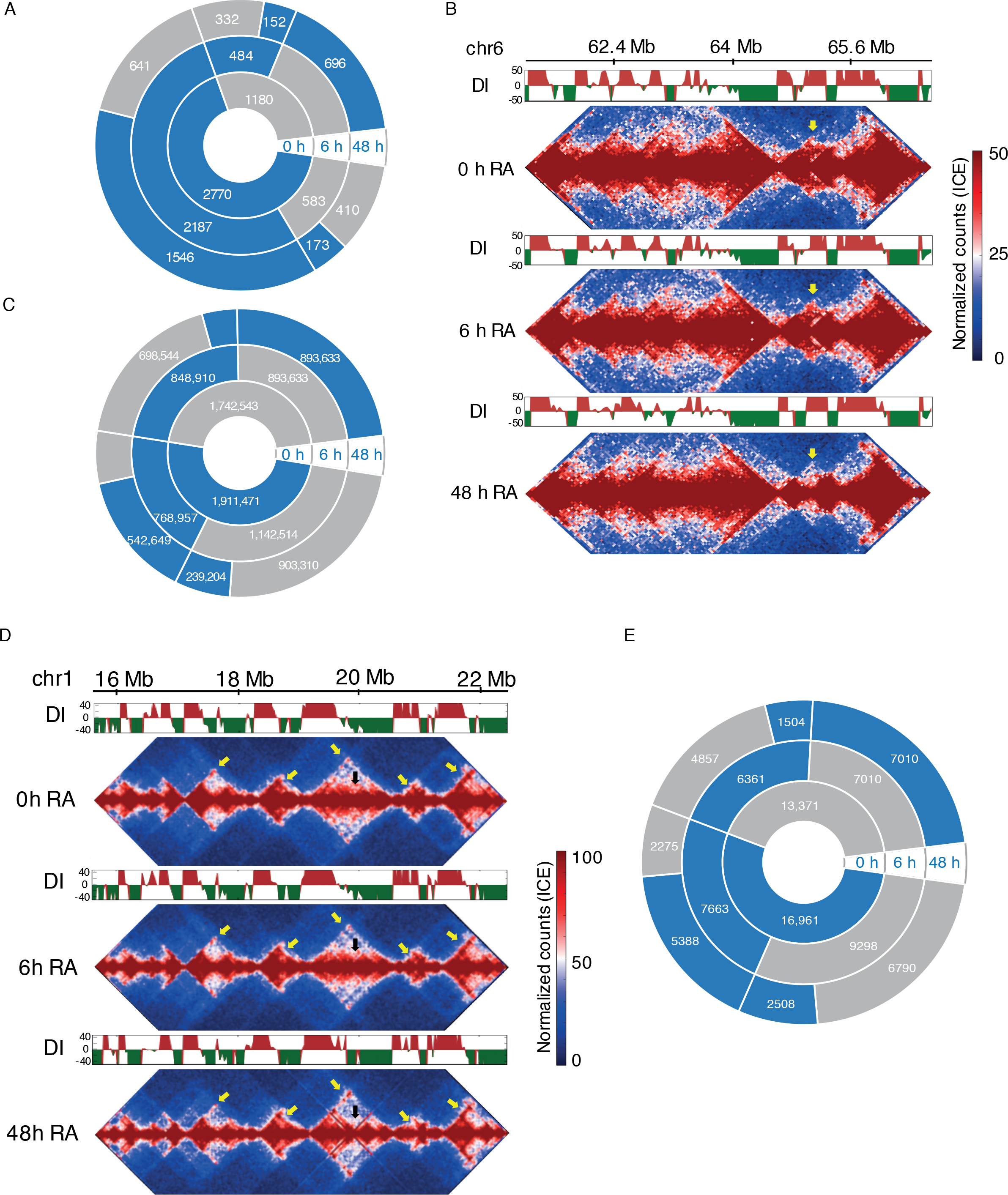
**Dynamics of chromatin organization** during the early stages of retinoic acid (RA) induced neuronal differentiation of P19 cells. **A.** TAD dynamics during the early stages of retinoic acid (RA) induced neuronal differentiation of P19 cells. The majority of TADs conserve their borders, however, several hundred were reorganized resulting in an increased number of unique TADs 48 h after morphogen treatment. Blue and grey - present and absent TADs at particular timepoint, respectively. **B.** Hi-C heatmaps with corresponding directionality indexes (DIs) displaying a TAD split at chromosome 6 (yellow arrows) after 48 h of treatment with RA. **C.** Genome-wide dynamics of long-range chromatin interactions. Blue and grey - present and absent interactions at particular timepoint, respectively. **D.** Hi-C heatmaps with corresponding directionality indexes (DIs) displaying the changes in the inner structure of TADs and overall loop dynamics (black and yellow arrows) after 6 h and 48 h of RA-induced neurogenesis. **E.** Dynamics of long-range chromatin loops connecting promoters of differentially expressed genes (DEGs) with their putative regulatory elements. Blue and grey - present and absent interactions at particular timepoint, respectively.

### Reconstruction and analysis of the GRN underlying early neural commitment informed by 3D chromosomal architecture

To identify key TFs and CREs driving the transcriptional changes during RA-induced neural commitment, we reconstructed a GRN based on the following information:

1. Data on differentially expressed genes (DEGs) in the course of RA-induced neural commitment in P19 cells, as well as the temporal profiles of RNA Pol II binding to identify actively transcribed genes (M.-A. Mendoza-Parra et al. 2016).
2. Promoter interaction data for the differentially expressed genes (DEG interactions) from the Hi-C analysis described above;
3. The temporal profiles of accessible chromatin regions detected by FAIRE-seq in the same system (M.-A. Mendoza-Parra et al. 2016) to identify putative active CREs;
4. The genome-wide binding sites of >600 TFs that we detected in >8,000 mouse ChIP-seq datasets deposited in the GEO database (that we made publicly available at http://ngs-qc.org/applications.php);
5. Transcription factor-target gene (TF-TG) relationships downloaded from the CellNet platform (Cahan et al. 2014; Morris et al. 2014).

We defined active CREs as accessible regions (based on FAIRE-seq) with the binding signals of TFs expressed at the same timepoint (see Methods for details). We then identified the putative target genes of active CREs based on Hi-C data, restricting the analysis to DEGs. This way, TFs were connected with their regulated genes through CREs and chromatin-mediated interactions. We supplemented these data with complementary information on the target genes of TFs from gene expression network analysis across multiple cell types and tissues available in the CellNet platform (Fig. 2A), which we again restricted to expressed TFs and DEGs. The nodes in the reconstructed GRN thus corresponded to active CREs, genes showing dynamic expression (DEGs) and regulator proteins (expressed TFs, some of which encoded by DEGs). In turn, the edges in the GRN reflected various types of spatiotemporal regulatory relationships between TFs, CREs and their target genes.

**Fig. 2.**
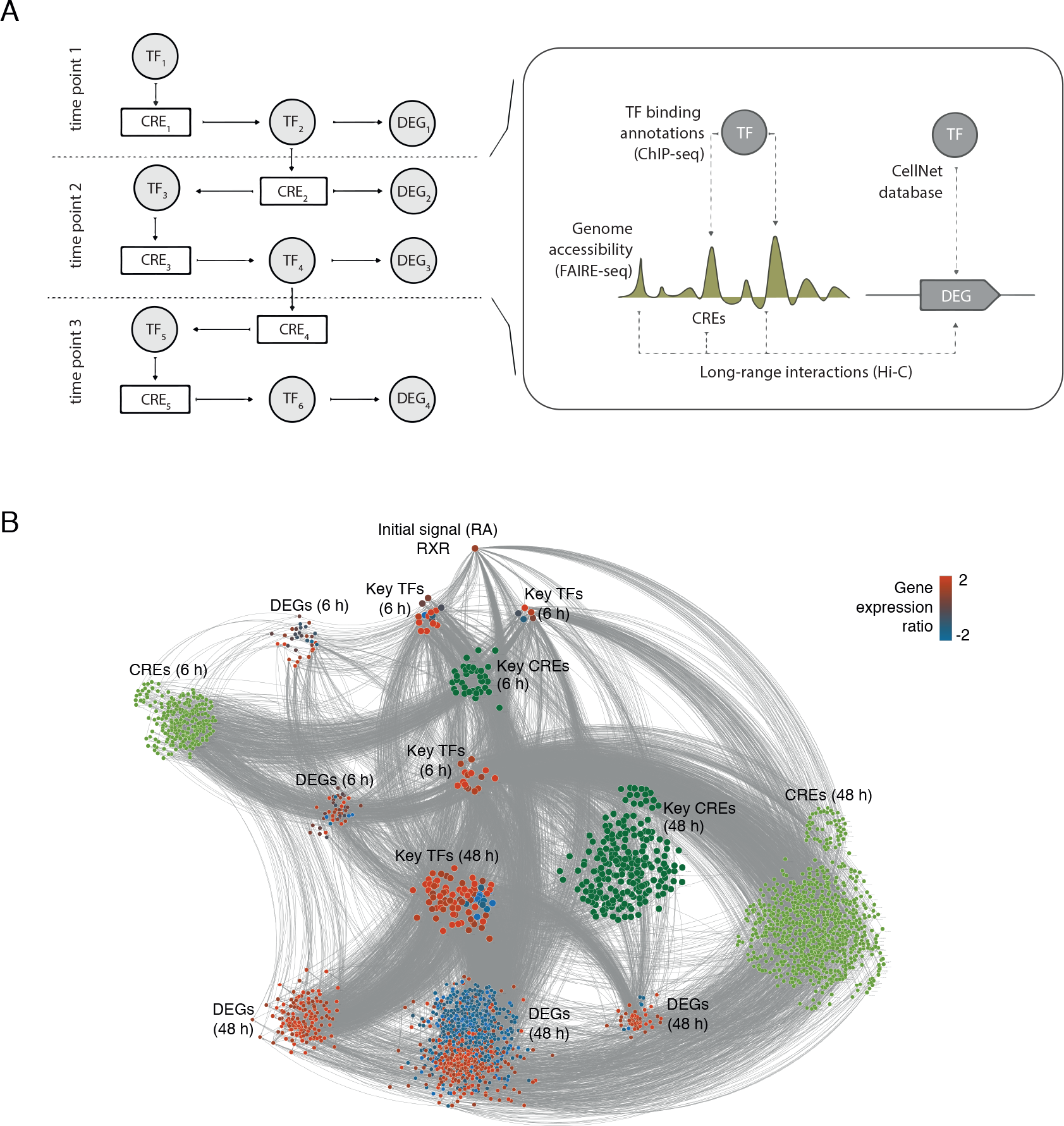
**Reconstructed Gene Regulatory Network** of early RA-induced neurogenesis. **A.** Scheme of data integration: transcriptome, CellNet TF-TG correlations, chromatin accessibility, TF-targeted ChIP-seqs and Hi-C long-range interactions. CRE, cis-regulatory element. **B.** Reconstructed GRN. Nodes represent genes (red and blue nodes corresponding to up- and downregulated genes, respectively, at 48 h of RA treatment) and cis-regulatory elements (light and dark green; key CREs - dark green). Node color shows the gene expression ratio at 48 h after RA treatment relative to non-differentiated sample as indicated. The size of nodes corresponds to the yield of signal propagation as predicted by TETRAMER.

To track signal propagation through the reconstructed GRN, we took advantage of our recently developed software tool, TETRAMER (Cholley et al. 2018; M.-A. Mendoza-Parra et al. 2016). In TETRAMER, the regulatory activity of the nodes (in our case, CREs and TFs) at each timepoint is discretized (−1, 0, 1), and the ability of every node to trigger the observed downstream changes in gene expression (“up”, “down”, “neutral”) is evaluated, taking into account signal propagation in both time and space (TF-CRE, TF-gene, CRE-gene). Nodes whose activity is inconsistent with the observed transcriptional changes are pruned from the network. This analysis produced a refined GRN underlying neural commitment comprised of 2,548 nodes (1,199 DEGs and 1,349 CREs) and 10,858 edges (Fig. 2B, Data S1).

### GRN-loop reveals driver TFs and *cis*-regulatory DNA elements

We applied a network permutation approach to delineate nodes, whose ability to induce the observed gene expression changes underlying neural commitment was higher than that expected at random, revealing 125 putative driver TFs and 246 driver CREs (Fig. 2B; Data S1). The identified driver TFs included known important neural regulators ASCL1, GBX2, MEIS1, PAX6, TAL2 and others (Data S1), generally validating the predictive power of our method (Achim et al. 2013; Huang et al. 2012; Kallur et al. 2008; Marcos et al. 2015; M.-A. Mendoza-Parra et al. 2016; Nakayama et al. 2013). Below we describe several examples of the identified subnetworks comprised of driver TFs and CREs predicted by our approach (Fig. 3).

**Fig. 3.**
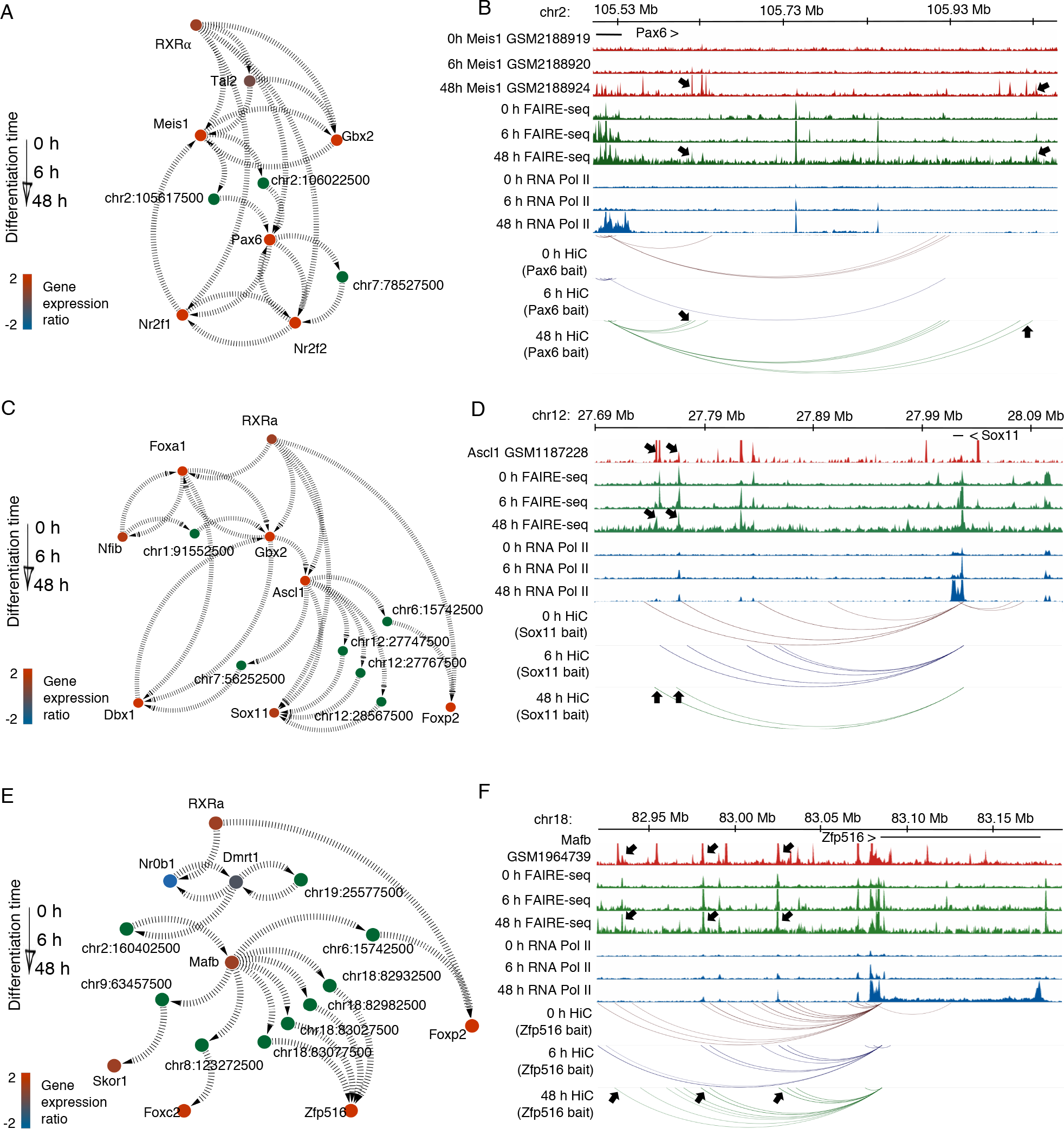
**GRN subnetworks** (**A**, **C** and **E**) propagating the initial driver signal – RA, activating RAR-RXRa - to the target nodes of the final phenotype (48 h) through the key regulatory TFs and key regulatory elements. The color of genes shows gene expression levels at 48 h after RA treatment; the color of enhancers reflects their “driver” or supportive role in signal propagation. Unidirectional arrows reveal the flux of the initial signal from the source node to the target node. Figures **B, D** and **F** give integrative views of the ChIP-seq profiles of the indicated TFs, the dynamics of chromatin accessibility (FAIRE-seq), RNA polymerase II recruitment (Pol II) and 3D chromatin organisation (Hi-C), and illustrates the state/occupancy of regulatory elements (black arrows) with key TFs regulating target gene expression and the signal propagation predicted by TETRAMER.

In the example shown in Fig 3A, RA induces TAL2 and GBX2 expression, and together they activate *Meis1*, which is upregulated in 48 h after RA treatment. At the same timepoint, MEIS1 binds two CREs on chromosome 2 (Figs. 3A, 3B; binding sites marked by arrows in 3B; Data S2), which become accessible at the same time point. Also at 48h, these CREs form a DNA looping interaction with the promoter region of *Pax6* (Fig. 3B; loops marked by arrows), consistent with the upregulation of PAX6 expression at this time point. We have further validated these interactions using 3C technique (Fig. S2A), and the regulation of *Pax6* by MEIS1 has been independently validated recently (Owa et al. 2018). The RA signal propagates further through PAX6 to activate interneuron-related genes *Nr2f2* and *Nr2f1* (Hu et al. 2017).

Fig. 3C shows a complex cascade of regulatory events predicted to underlie RA-induced activation of multiple neural-specific genes. RA treatment leads to transcriptional activation of ASCL1, which binds to a range of regulatory elements (Fig. 3D) on different chromosomes (Wapinski et al. 2013), which in turn propagate the signal to *Dbx1*, *Sox11* and *Foxp2* involved in neuronal fate specification (Bonev et al. 2017; Inamata and Shirasaki 2014; Karaz et al. 2016; Tsui et al. 2013; Wang et al. 2013) through DNA looping interactions (Figs. 3C, 3D; Data S3). We have also validated these interactions using 3C (Fig. S2B).

Finally, we have previously predicted and validated DMRT1 as a key factor involved in RA-induced neurogenesis (M.-A. Mendoza-Parra et al. 2016). Here, we show that this effect of DMRT1 is likely mediated, at least in part, by its binding to a CRE on chromosome 2 that forms a loop with *Mafb* (Figs. 3E, 3F; Data S4; see Fig. S2C for validation by 3C). MAFB, in turn, acts on several regulatory elements across the genome, which induce the expression of a number of neural TFs. Notably, unlike in the first example (Fig. 3A), the subnetworks in Fig. 3B and 3C included temporally invariant CRE-gene loops. Taken together, these results demonstrate how both pre-established and *de novo* created chromatin interactions participate in transcriptional regulation in early development, consistent with previous findings (Ghavi-Helm et al. 2014; Freire-Pritchett et al. 2017).

### Chromatin structure directs and constraints cell fate specification and lineage commitment

While the P19 ES-like cell line undergoes neural commitment in response to RA, a related cell line, F9 differentiates towards the endodermal lineage under the same conditions (REF). Strikingly, we previously showed that the two cell lines exhibit markedly similar profiles of gene expression and chromatin state (M. A. Mendoza-Parra et al. 2014; M.-A. Mendoza-Parra et al. 2016). We therefore hypothesised that their differential response to the same morphogen might be underpinned by different chromosomal organisation. To verify this, we performed Hi-C in RA-treated F9 cells at the same time points (0, 6, 48h) as for P19 in the analysis above. Consistent with our hypothesis, the two cell lines showed differences in higher-order chromatin architecture, as exemplified by only 1008 shared TADs (within 80 kb window for each border), corresponding to 36% of P19 TADs and 47% of F9 TADs (Figs. 4A, 4B).

**Fig. 4.**
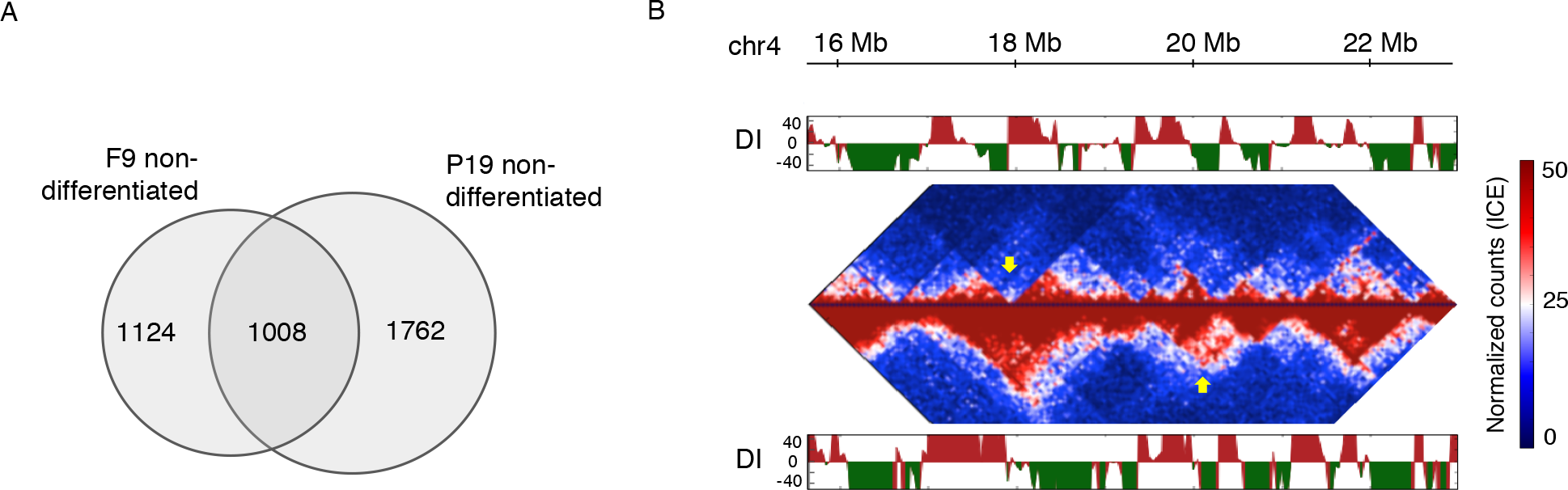
Cross-cell fate comparison of chromatin organization. **A.** Comparison of global chromatin organisation between F9 and P19 at non-differentiated state. TAD border comparison shows large differences in higher-order chromatin structure. **B.** Example of TAD and inner TAD structure differences in non-differentiated cell lineages.

We then detected promoter-based DNA loops in F9 Hi-C data and asked which of the driver CRE-target gene connections were shared between P19 and F9 cells during the first 48h of treatment and which ones were specific to P19 cells. In total over all time points, we identified 470,255 shared and 3,183,759 P19-specific interactions. Examples of shared CRE-gene connections include those between CREs and *Sox1* and *Zfp516* genes in Figures 3C and 3E. In contrast, we found that P19 driver CRE connections of Pax6 (Fig. 3A, Data S2) were absent in F9 during the entire period of RA treatment (Fig. S3). These striking differences in the chromatin interactions within the RA-Gbx2-Pax6 subnetwork between the two cell lines might contribute to their differential cell fate in response to RA. Jointly, our observations support the role of chromatin organisation in lineage choice and highlight the role of chromatin dynamics in the propagation of transcriptional regulatory signals in the course of lineage commitment.

## Discussion

In this study, we presented a network reconstruction and analysis approach - GRN-loop - integrating the transcriptome, epigenome and chromatin interaction dynamics. Applying GRN-loop to a model of morphogen-induced early neuronal lineage commitment, we revealed a set of novel and previously known driver TFs and CREs. We show how signal propagation through the GRN depends on a combination of pre-established and de novo-generated connections between CREs and target genes during lineage commitment, echoing the global reorganisation of chromosomal interactomes in this process (Freire-Pritchett et al. 2017). Finally, by comparing the 3D interactomes of two ES-like cell lines showing differential lineage choice in response to the same morphogen, we demonstrate how cell line-specific chromatin interactions between CREs and their target genes likely play a key role in this phenomenon.

GRNs are a powerful tool to study developmental processes (reviewed in (Peter and Davidson 2017). While many GRN approaches rely on co-expression data to reconstruct the network, epigenetic data provide promising opportunities for enhanced GRN reconstruction (Ramirez et al. 2017; Rendeiro et al. 2016). For example, a recent study used chromatin accessibility information to improve the inference of TF-target gene associations (Miraldi et al. 2019). Network propagation in reconstructed GRNs offers a way to simultaneously consider all possible paths between genes, and prioritise genes and modules based on their connectivity and prior information from multiple data sets (reviewed in (Cowen et al. 2017)). This approach has been successfully used for gene function and drug target prediction (Ideker and Sharan 2008; Csermely et al. 2013; Sharan, Ulitsky, and Shamir 2007) and disease characterization (Ideker and Sharan 2008). In our previous work, we have applied signal propagation to identify TFs driving the cell fate specification programs, using genome-wide gene expression data, as well as chromatin accessibility information to define a subset of proximal TF-gene associations (M.-A. Mendoza-Parra et al. 2016). Here, we have significantly enhanced this approach by incorporating data on 3D chromatin organisation and chromatin accessibility globally into the network, thus accounting for TF binding at CREs and their regulatory effects on target genes.

Our study has capitalised on ES-like cell lines owing to their notable property of showing a markedly different and homogeneous developmental response to the same morphogen and the wealth of developmental course data that we have previously generated in this system. However, the conceptual and methodological framework of GRN-loop can be readily applied to other cell types including ES cells, for which information on chromatin accessibility and chromosomal is available from any technique (such as ATAC-seq, DNAse-seq; and Hi-C, HiChIP, CHiA-PET, Capture Hi-C, respectively), to provide an integrated understanding of dynamic biological processes.

## Methods

### Cell culture

F9 cells were cultured in Dulbecco’s modified Eagle’s medium (DMEM) supplemented with 10% fetal calf serum (FCS) and 4.5 g/l glucose; P19 cells were grown in DMEM supplemented with 1 g/l glucose, 5% FCS and 5% delipidated FCS. Both media contained 40 μg/ml Gentamicin. F9 or P19 EC cells were cultured in monolayer on gelatine-coated culture plates (0.1%). For cell differentiation assays, RA was added to plates to a final concentration of 1μM for different exposure times.

### Transcriptome and Epigenome assays

We have previously described the data on transcriptome dynamics and chromatin immunoprecipitation assays (M.-A. Mendoza-Parra et al. 2016); the datasets are available from the Gene Expression Omnibus (GEO) database (GSE68291). ChIP-seq profiles for MEIS1, ASCL1 and MAFB are accessible from GEO (GSM2188919, GSM2188920, GSM2188924, GSM1187228 and GSM1964739).

### Hi-C experiments

Chromatin organization has been assessed by Hi-C (Lieberman-Aiden et al. 2009). The original Hi-C protocol has been improved, increasing the ligation yields and modifying the steps that favour chromatin de-crosslinking, while keeping the conventional Hi-C workflow. Briefly, cells were crosslinked with 1% PFA (Electron Microscopy Sciences) for 30 min at room temperature, after which cells were washed twice in ice-cold PBS. Cells were collected by centrifugation (1000 r.p.m. for 10 min at 4°C), washed once with PBS followed by a second centrifugation step. The supernatant was removed and cell pellets were snap frozen in liquid nitrogen and stored at −80°C. Per Hi-C sample 20-25M cells were used. Cell pellets were incubated in 1 ml ice-cold lysis buffer (10 mM Tris-HCl pH 8, 10 mM NaCl, 0.2% NP-40, protease inhibitor cocktail (Roche) for 15 min on ice. The cells were Dounce homogenized (30 strokes, 1 min pause on ice, 30 strokes). After centrifugation (2500 r.p.m. for 5 min at 4°C), nuclei were washed once with NEBuffer 2 and resuspended in 250 ml of NEBuffer 2. The sample was split into aliquots of 50 µl. Each sample was further complemented with 307 µl of NEBuffer2, 38 µl of 1% SDS (0.1% final concentration), incubated for 10 min at 65°C and put on ice directly afterwards. To quench SDS, 44 µl of 10% Triton X-100 was added. The restriction digest was set up directly afterwards and performed overnight at 37°C with agitation (750 r.p.m.) with *HindIII* (NEB; 20 µl of 20 U/µl). Using biotin-14-dCTP (Life Technologies), dATP, dGTP and dTTP (all at a final concentration of 28 µM), the *HindIII* restriction sites were then filled in with Klenow (Thermo Fisher Scientific) for 75 min at 37°C mixing at 750 r.p.m. for 5 s each 5 min. 3C sample was generated from 5M cell aliquots in parallel with Hi-C by replacing the fill-in mix with corresponding amount of nuclease-free water and treated in same way as Hi-C sample. The fill-in reaction was directly followed by overnight ligation at 16°C in a total volume of 8.2 ml ligation buffer (50 mM Tris-HCl, 10 mM MgCl_2_, 1 mM ATP, 10 mM DTT, 100 µg/ml BSA, 1% Triton X-100, 10,000 units T4 DNA ligase (NEB 400U/µl)) per 5 million cells starting material. After ligation, reverse crosslinking (65°C overnight in the presence of Proteinase K (Roche)) was followed by two sequential phenol/chloroform/isoamylalcohol extractions. Purified solution was then concentrated on 30K AMICON 15 ml centrifugal filter (span at 3000 r.p.m.) to 300 µl and DNA was precipitated in ethanol for 1 hour at −80°C. The DNA was span down (13,000 r.p.m. for 30 min at 4°C). The pellets were washed twice in 70% ethanol, dried and resuspended in 100 µl TE buffer. Hi-C samples were combined and washed twice on Amicon 0.5 ml centrifugal filter unit with TE and concentrated to 100 µl in case of Hi-C sample and to 25-30 µl in case of 3C sample. DNA concentration was determined using a Nanodrop. The efficiency of biotin incorporation was assayed by amplifying a ligation product, followed by digestion with *HindIII* or *NheI*. If the ligation product was fully or almost fully digested (Fig. S4) by NheI, Hi-C sample was used further.

To remove biotin from non-ligated fragment ends, 40 µg of Hi-C library DNA were incubated with T4 DNA polymerase (NEB) for 4 hours at 20°C, followed by phenol/chloroform purification and DNA precipitation overnight at −20°C or for 1 hour at −80°C. After DNA pellet resuspension, the sonication was carried out to generate DNA fragments with a size peak around 200 bp (Covaris Sonolab 7 settings: duty factor: 10%; peak incident power: 175W; cycles per burst: 200; time: 60 sec).

A double size selection using AMPure XP beads (Beckman Coulter) was performed to select 100-400 bp DNA fraction. The ratio of AMPure XP beads solution volume to DNA sample volume was adjusted to 0.7:1. After incubation for 10 min at room temperature, the sample was transferred to a magnetic separator (DynaMag-2 magnet; Life Technologies), and the supernatant was transferred to a new 1.5 ml tube, while the beads were discarded. The ratio of AMPure XP beads solution volume to DNA sample volume was then adjusted to 1.1:1 final. After incubation for 15 min at room temperature, the sample was transferred to a magnet (DynaMag-2 magnet; Life Technologies) and washed twice with 80% ethanol. The DNA was eluted in 100 µl of TLE (10 mM Tris-HCl pH 8.0; 0.1 mM EDTA).

Biotinylated ligation products were isolated using MyOne Streptavidin C1 Dynabeads (Life Technologies) on a DynaMag-2 magnet (Life Technologies) in binding buffer (5 mM Tris pH8, 0.5 mM EDTA, 1 M NaCl) for 45 min at room temperature. After one wash in binding buffer and two washes in wash buffer (5 mM Tris, 0.5 mM EDTA, 1 M NaCl, 0.05% Tween-20) the DNA-bound beads were resuspended in a final volume of 55.5 µl of nuclease-free water and used for end repair, adaptor ligation and PCR amplification with NebNext Ultra library preparation kit for Illumina (NEB #E7370S) following the manufacture protocol. Note that after adaptor ligation, beads were washed twice in washing buffer, twice in 10 mM Tris-HCl pH 8.0, resuspended in 25 µl of 10 mM Tris-HCl pH 8.0 followed by PCR. For the PCR amplification 5 µl DNA-bound beads were used with Index primer and universal PCR primer diluted 1/10. Fve PCR cycles were made to obtain the Hi-C library that was purified with AMPure XP beads at 1:1 ratio, followed by paired-end high-throughput sequencing on HiSeq4000.

### 3C validation of PIR – promoter interactions

3C-PCR validation has been performed on 3C samples produced in parallel with the corresponding Hi-C samples. Corresponding primers are provided in Data S5.

### Hi-C data processing

Raw paired-end sequencing reads were mapped against the mouse genome (mm9) with Bowtie2. Low quality reads (MAPQ<=10), PCR duplicates and interactions falling within the same restriction fragment were filtered out. Hi-C contact maps were constructed at 5 kb resolution, then normalized by matrix balancing using the ICE algorithm. Importantly, to avoid the contamination of the downstream analysis by over-normalized interactions, the cut-off for the extremely low and high occupancy bins has been adjusted individually for each sample, instead of using fixed thresholds proposed in the ICE method. To define the upper and lower outliers cut-off, we evaluated the distribution of total counts per bin and applied the modified z-score (outliers > 3.5) to fix the upper cut-off and the first local minima to define the lower cut-off (Fig. S5). Statistically significant interactions were identified by Fit-Hi-C (Ay, Bailey, and Noble 2014). To define reproducible interactions between biological replicates we used sdef (Blangiardo, Cassese, and Richardson 2010) R package. Only interactions that appear in both biological replicates, pass the cut-off defined by sdef and have a normalized count greater than or equal to 3 in both replicates were considered as significant and have been used for further integrative analysis and GRN reconstruction (Data S6).

### Chromatin structure, epigenome and transcriptome integration

We have associated open-chromatin regions - defined by FAIRE-seq assay - to the promoter-associated distal GAPs. FAIRE localization sites were then compared with a comprehensive collection of TF ChIP-seq assays retrieved from the public domain (M.-A. Mendoza-Parra et al. 2013). Note that the TF collection in use in this study includes a large number of datasets in addition to those provided by the ENCODE consortium; thus, our analysis is a comprehensive comparative study not only because of the large number of datasets used but also with respect to the diversity of the cellular systems considered. Transcriptome, RXR binding sites from ChIP-seq, TF annotations from public datasets and Hi-C long-range chromatin interactions were integrated and visualized using the Cytoscape platform (version 3.7.1). The signal propagation was performed multiple times using TETRAMER tool and a randomized network approach as control (Cholley et al. 2018).

## Acknowledgments

We thank Peter Fraser and William Orchard for the critical review of the manuscript and helpful comments. We thank the IGBMC Microarray and Sequencing platform (France Génomique consortium—ANR-10-INBS-0009). This study was supported by AVIESAN-ITMO Cancer, the Ligue National Contre le Cancer, the Institut National du Cancer (INCa) and the Agence Nationale de la Recherche (ANRT-07-PCVI-0031-01, ANR-10-LABX-0030-INRT and ANR-10-IDEX-0002-02).

## Author contributions

H.G. designed the study; V.M. performed the experiments; V.M., M.A.M.P., M.B., M.S. and H.G. analyzed the data; V.M., M.S. and H.G. wrote the manuscript.

## Corresponding authors

Correspondence to Valeriya Malysheva or Hinrich Gronemeyer.

## Data and materials availability

Sequencing data reported in the current study were deposited at the Gene Expression Omnibus repository (GEO) of the National Center for Biotechnology Information under the GEO accession number GSE111875.

