## Supplemental Figure 1 for "Gene regulatory network reconstruction incorporating 3D chromosomal architecture reveals key transcription factors and DNA elements driving neural lineage commitment"

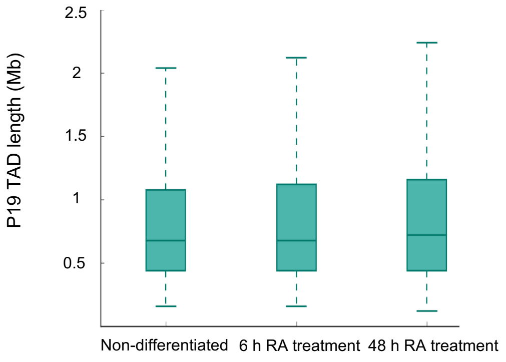


**Fig. S1. TAD and chromatin loops length in early RA-induced neurogenesis.** TAD size is largely conserved during early neurogenesis.
