## Supplemental Figure 2 for "Gene regulatory network reconstruction incorporating 3D chromosomal architecture reveals key transcription factors and DNA elements driving neural lineage commitment"

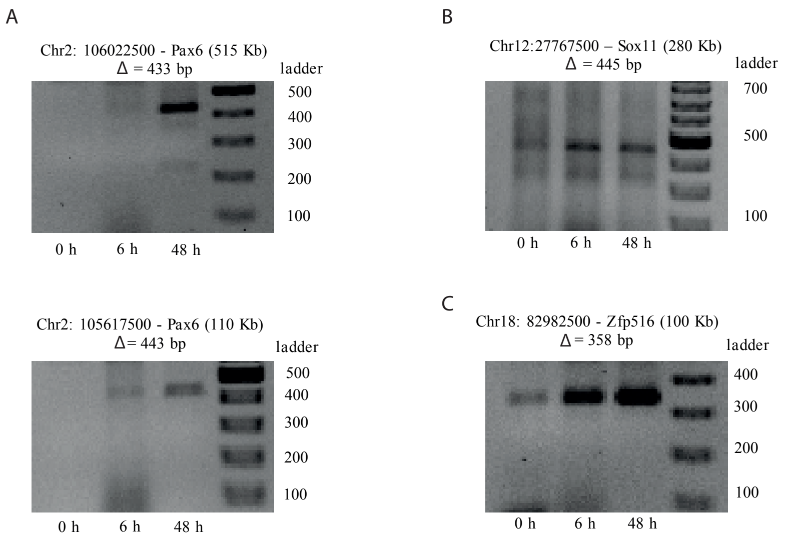


Fig. S2. 3C validation of PIR – promoter interactions. 3C-PCR validation of several long-range chromatin interactions predicted as key for the signal propagation, represented in Fig. 3 of the main text. Corresponding primers are provided in Data S.
