## Supplemental Figure 3 for "Gene regulatory network reconstruction incorporating 3D chromosomal architecture reveals key transcription factors and DNA elements driving neural lineage commitment"

Fig. S3

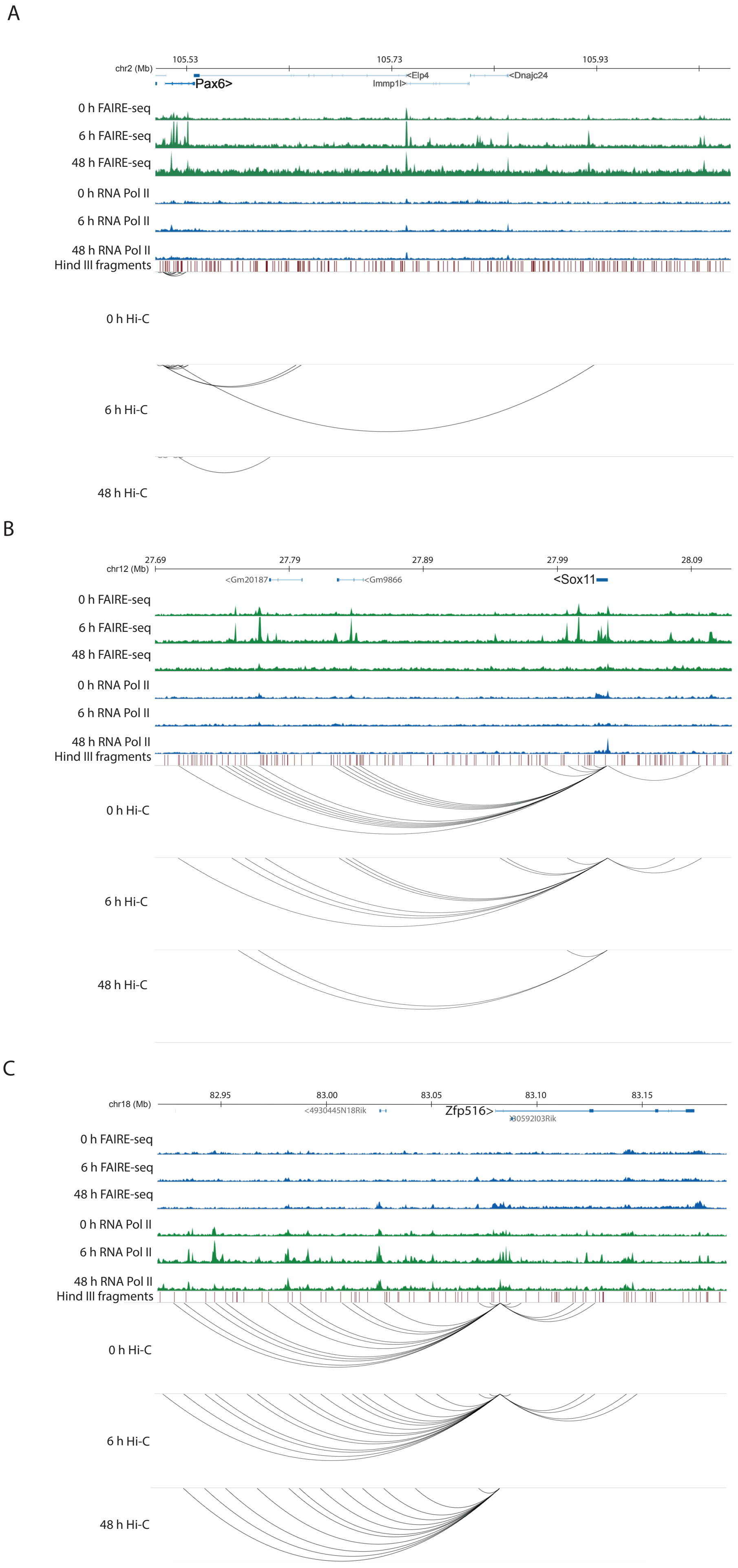


**Fig. S3.** Integrative views of the chromatin accessibility dynamics(FAIRE-seq), RNA polymerase II recruitment (Pol II) and 3D chromatin organisation (Hi-C) provide examples of A. cell-type specific chromatin interactions (CRE connections of Pax6, Fig. 3B) and B. and C. shared between F9 and P19 chromatin interactions (between CREs and Sox1 and Zfp516 genes, Fig. 3C and 3E).
