## Supplemental Figure 4 for "Gene regulatory network reconstruction incorporating 3D chromosomal architecture reveals key transcription factors and DNA elements driving neural lineage commitment"

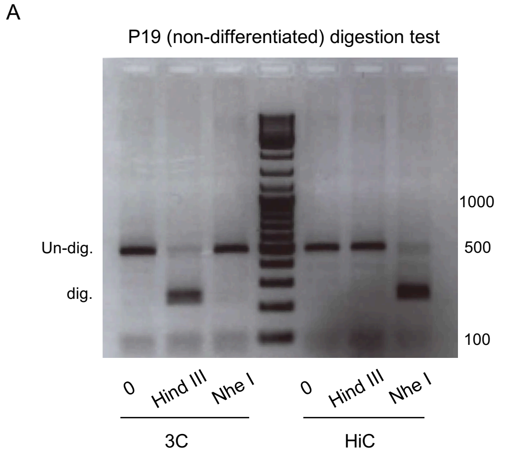


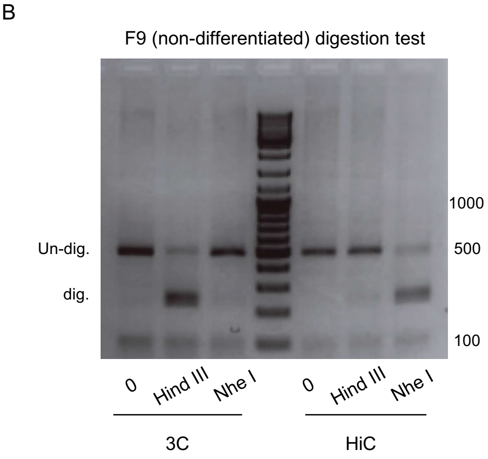


**Fig. S4. HiC and 3C digestion tests**. Digestion of a PCR amplicon generated from non-neighboring restriction fragments in both 3C and Hi-C sample. The amplicon was digested with *HindIII*, *NheI*, or not digested (0). High digestion rates with the *NheI* in the HiC sample indicates high filling-in and ligation efficiency. Representative cases for P19 (**A**) and F9 (**B**) samples are shown.
