## Supplemental Figure 5 for "Gene regulatory network reconstruction incorporating 3D chromosomal architecture reveals key transcription factors and DNA elements driving neural lineage commitment"

Fig. S5

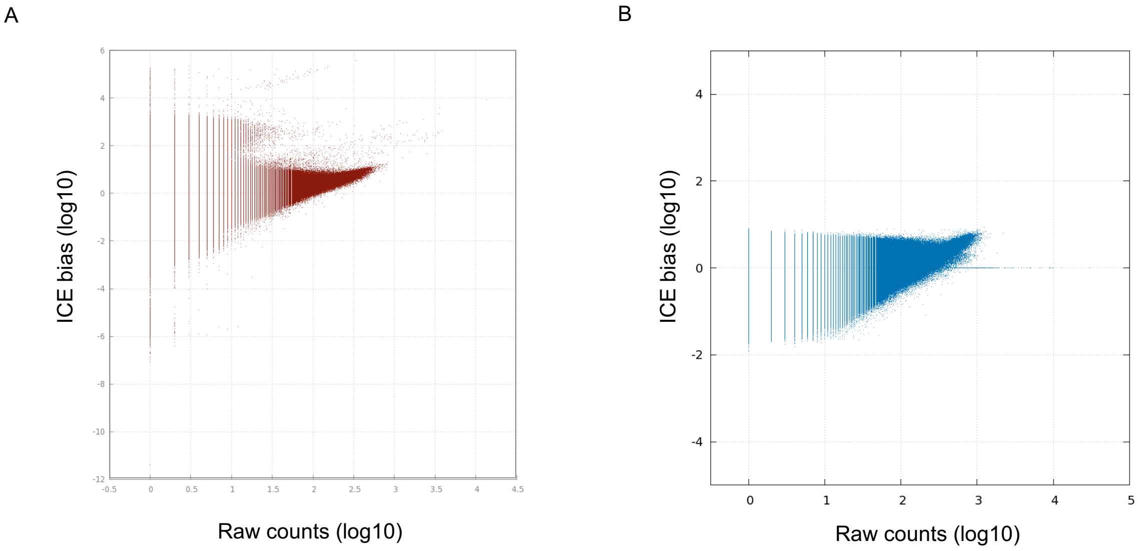


**Fig. S5.**
**HiC normalization adjustment.** ICE bias vs raw count distribution when applying **A.** the standard ICE thresholds and **B.** sample-specific thresholds, showing that low count bins are not transformed into bins with artificially high count, enabling further downstream analysis.
